# Citizen science and aquatic macroinvertebrates: public engagement for catchment-scale pollution vigilance

**DOI:** 10.1101/842559

**Authors:** Adam Moolna, Mike Duddy, Ben Fitch, Keith White

## Abstract

Citizen science aims to engage the wider population beyond scientists and statutory agencies, providing a catalyst for positive change and influencing policymakers and institutions. The Riverfly Partnership has been supporting a growing network of citizen science volunteers monitoring aquatic macroinvertebrates in rivers and streams across the United Kingdom since 2004. In Manchester, Salford and surrounding areas, Riverfly Partnership monitoring began in 2011 with volunteers from a catchment-wide fishing club. This provided a catalyst for broader public engagement, wider environmental projects, and the establishment of a new grassroots environmental charity. The vigilance of the network demonstrated its value by flagging a major pollution event wiping out all macroinvertebrates along 19 km of the River Irwell in April 2017. By evaluating monitoring data and the citizen science program’s impacts, we identify key lessons for taking forward public engagement in river catchment management both in Greater Manchester and elsewhere.

## 1 INTRODUCTION

### 1.1 Macroinvertebrates as bioindicators

The macroinvertebrates living amongst vegetation and within or on the sediment of rivers provide a valuable resource for the assessment of a wide range of aquatic parameters (Armitage et al. 1983; Hawkes 1998; Everard 2008). With the long residence time of individuals (typically one to three years), the macroinvertebrate community acts in effect as a data logger: its composition indicates the long-term average of water quality and whether the site experiences occasional intense pollution episodes. Macroinvertebrates can also indicate the flow regime and habitat and substrate parameters (Extence et al. 1999; Glendell et al. 2014).

Biological monitoring of rivers with macroinvertebrates is used worldwide from Europe and the Americas to Africa and Australasia (Smith et al. 1999; Bonada et al. 2006; Ollis et al. 2006; Shull & Lookenbill 2007; Birk et al. 2012). Rigorous statutory monitoring was developed in the UK through the 1970 National River Pollution Survey (Hawkes 1998) and subsequent BMWP (Biological Monitoring Working Party) system, which scores taxa on a scale 1-10 based on their tolerance to pollution (Armitage et al. 1983). Each taxon identified in a sample is scored and the BMWP score is the total of the taxa scores. A further development was Average Score Per Taxon (ASPT), which is the BMWP cumulative score divided by the number of taxa and hence a measure of the average pollution tolerance of the sample (Everall et al. 2017). The ASPT limits both the influence of sampling effort (larger samples are likely to include more families, which increases the BMWP score) and the skewing effect of an outlier.

The macroinvertebrate sampling technique used in statutory monitoring in the UK is that of a semi-quantitative kick sample (UKTAG 2008). A standardized net catches macroinvertebrates dislodged by an operative disturbing the substrate with the feet for a fixed period of time, taking a zigzag route across the river to ensure that all habitats are included (Furse et al. 1981; Armitage et al. 1983). The same sampling technique can be employed with minimal training by the general public and is therefore readily used in citizen science monitoring programs. Assessment of samples by BMWP or ASPT scoring, however, requires a professional level of taxonomic expertise to identify 84 families of macroinvertebrates plus the Class Oligochaeta (Hawkes 1998).

### 1.2 Riverfly Partnership citizen science

Macroinvertebrates are of particular interest to anglers because of their influence on fish distributions (Wallace & Webster 1996) and the use of macroinvertebrate imitations (“flies”) to catch fish such as salmon and trout. The Riverfly Partnership identified anglers as the focus constituency for citizen science monitoring of river macroinvertebrates when it began in 2004 with collaboration between the Freshwater Biological Association, the Natural History Museum London, the Salmon & Trout Association (now known as Salmon & Trout Conservation UK) and other interested parties.

The Riverfly Partnership’s Anglers’ Riverfly Monitoring Initiative (ARMI) index uses a selection of the moderately tolerant to pollution sensitive BMWP families (BMWP values 4-10) placed into 8 groups for simplified identification in a live sample (Riverfly Partnership, http://www.riverflies.org; Di Fiore & Fitch 2016). The ARMI index does not distinguish between pollution tolerances of the 8 groups (each is treated equivalently in the scoring formula) but does place a weighting on the abundance of each group, estimated as logarithmic categories. The ARMI score is the total of the abundance scores for the eight groups: zero individuals of a group scores 0; 1-9 scores 1; 10-99 scores 2; 100-999 scores 3; and 1000 or more scores 4. The simplified target groups, live sorting and broad abundance categories make riverside sampling and processing accessible to non-scientist trainees with a one day training course. Sample processing can be on the riverside: identification is straightforward; equipment requirements are simple; no preservatives are needed; and estimate of numbers is quicker and easier than counts to the last individual.

### 1.3 The context of Greater Manchester’s rivers

Manchester, recognized as the world’s first industrial city, has a long history of polluted rivers (Burton 2003). The influence of its industrial past is still seen today with continuing leachate of heavy metal contaminants into the River Irwell (Hurley et al. 2017), which is 63 km long and drains a catchment of 701 km^2^ including much of Greater Manchester. Water quality improvements during the 20^th^ Century along the River Irwell and the River Mersey into which it flows downriver of Manchester were boosted from 1985 by the UK government initiated Mersey Basin Campaign (Gregory et al. 2011).

However, continuing episodic pollution events of raw sewage released by combined sewer overflows following heavy rainfall plus industrial discharges remain an ongoing concern of local recreational anglers in close contact with the rivers in and around the Greater Manchester conurbation. Incidences of fish death suspected to be connected with sewage and industrial discharges led to the Salford Friendly Anglers’ Society (SFAS) joining the Riverfly Partnership’s citizen science program in September 2011.

The SFAS was founded in April 1817 and is an example of the “friendly societies” of 18^th^ and 19^th^ Century Britain, which were social and insurance clubs usually for people sharing common interests such as angling (Cordery 1995; Gorsky 1998). Legislative changes in the 1890s meant the financial role ceased but the SFAS continued as a fishing club; it is today an organized group of anglers focused on fishing the River Irwell and its tributaries. Free membership, free fishing and other free activities are a central tenet of the SFAS to maximize engagement and ensure accessibility for social deprived communities around the urban stretches of the river.

### 1.4 This study

There has been a growing consensus that citizen science monitoring can make important contributions to management aimed at improving water quality and in responses to pollution incidents (Latimore & Steen 2014; Shupe 2016; Shuker et al. 2017).

This study tests the hypothesis that citizen science can both inform and catalyze river improvements by assessing the Riverfly Partnership citizen science program in and around the urban conurbation of Greater Manchester. The vigilance of the network and its responsiveness is demonstrated with volunteer-led identification of the location of a pollution event causing a large-scale macroinvertebrate kill along 19 km of the River Irwell in April 2017.

The capacity for citizen science engagement of communities to catalyze river improvements is demonstrated by restoration and rehabilitation projects that have emerged alongside and by the formation of a new grassroots environmental charity. We examine the challenges the program has faced and continues to face, the lessons learnt, and recommendations to guide future citizen science work monitoring and reporting on river health in Manchester and beyond.

## 2 METHODS

### 2.1 Macroinvertebrate sampling, biotic indices and laboratory methods

Macroinvertebrate kick samples were taken for a cumulative three minutes along a 10-20 m length of river, sampling habitats and substrates proportional to their abundance, plus a one minute search of larger stones for attached macroinvertebrates. Riverfly Partnership ARMI scoring (Riverfly Partnership, http://www.riverflies.org; Di Fiore & Fitch 2016) recorded 8 groups of macroinvertebrates: cased caddis (Trichoptera); caseless caddis (Trichoptera); true mayfly (Ephemeridae); flat-bodied mayfly (Heptageniidae); blue-winged olives (Ephemerellidae); olives (Baetidae); stoneflies (Plecoptera); and freshwater shrimp (*Gammarus* spp.). BMWP and ASPT scoring (Whalley & Hawkes 1996) recorded all macroinvertebrates, except oligochaetes, to family level using Croft (1986) and Mellanby (1963). Autumn was defined for data handling following UKTAG (2014) as 1^st^ September to 30^th^ November.

### 2.2 Sampling sites

Eleven sites along the River Irwell from Rawtenstall in the upper reaches (13.5 km from source) to Strangeways adjacent to Manchester city centre (56.5 km from source) and two additional sites on the River Roch and Eagley Brook tributaries were selected to cover a range of putative clean and polluted waters and of semi-rural to urban locations across the Irwell catchment. Sites were sampled monthly in September, October and November 2011 to establish baseline Riverfly Partnership ARMI scores. For comparison of the ARMI index with BMWP and ASPT indices, 8 of the Riverfly Partnership reference sites and one additional site between Springwater Park and Burrs Country Park were sampled in January and March 2012. Sampling continued for nearly 4 years on the River Roch and continues to the present (6 years of data) at Ewood Bridge on the River Irwell and at the Eagley Brook tributary. Additional sites were sampled in April 2017 between the Chatterton Country Park and Ewood Bridge reference sites in response to a suspected pollution event to constrain the upriver location at which the macroinvertebrate kill began.

### 2.3 Mapping and data analysis

Maps were generated and distances along the river measured using ArcMap and QGIS software. Statistical analysis and graphs were produced using Minitab and Microsoft Excel software. Correlations of ARMI scores with BMWP or ASPT scores were tested using Spearman’s correlation coefficient.

## 3. RESULTS

### 3.1 Sites, baseline scores and long-term records

For 11 sites along the River Irwell and two additional sites on the Eagley Brook and River Roch tributaries (Fig. 1) an autumn (September to November) baseline of ARMI scores was established (Fig. 2). The ARMI baseline of 5.7 at Rawtenstall (13.4 km from the source) increases to 9.7 at Springwater Park 22.2 km downriver; then decreases to 5.3 at Sion Street 3.9 km below Springwater Park. The baseline falls as the Irwell flows through the urban conurbation towards the city center, with low scores of 2.7, 2.0 and 2.0 at Clifton Country Park, Agecroft and Strangeways, respectively (14.4 km, 8.7 km and 0.6 km from the edge of the city center, respectively). Autumn baseline ARMI scores for sites on Irwell tributaries were 7.7 for Eagley Brook and 6.5 for the River Roch (8.7 and 12.0 km from the boundary of the city center, respectively).

**Fig. 1.**
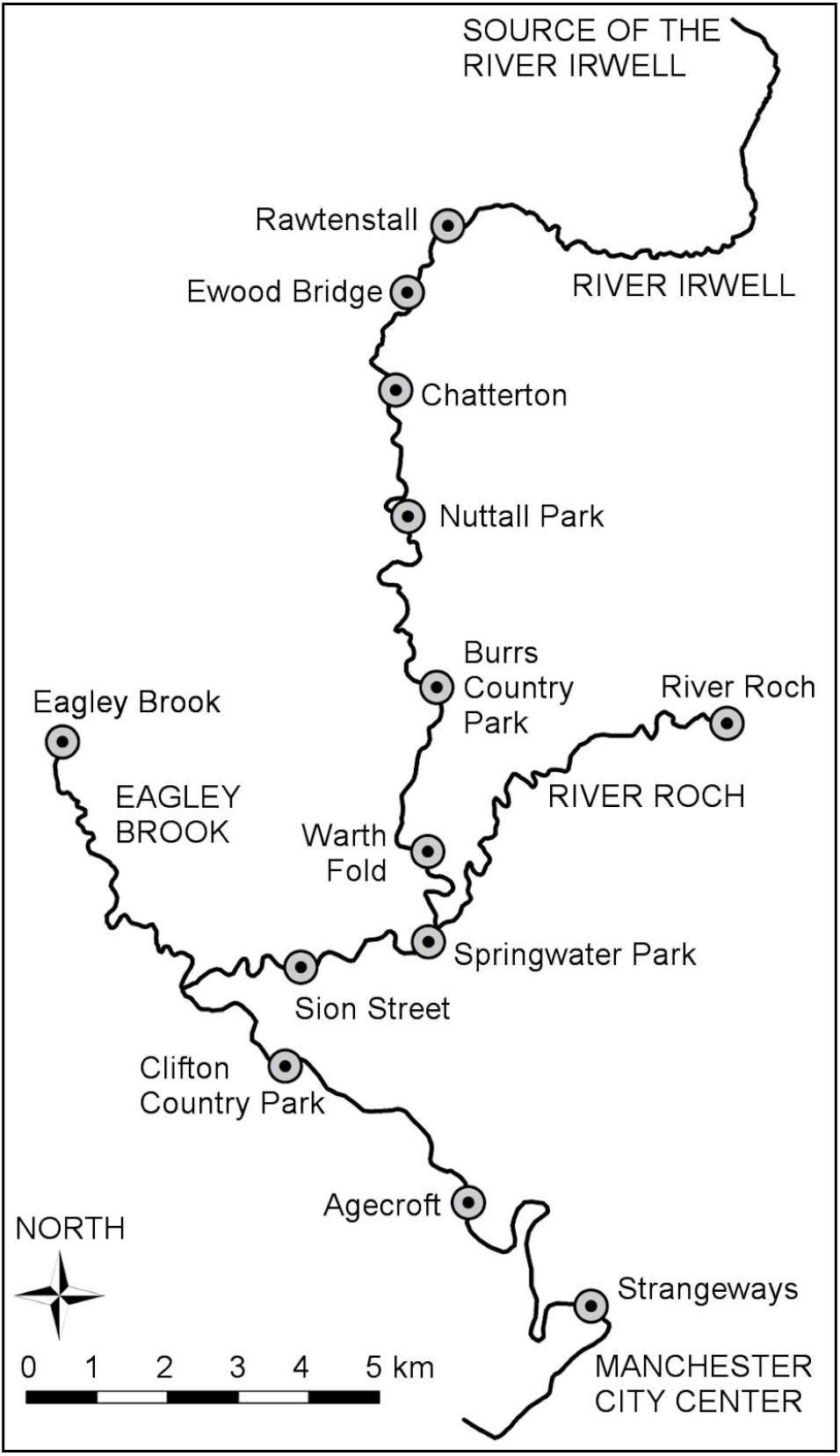
Riverfly Partnership reference sites on the River Irwell and tributaries

**Fig. 2.**
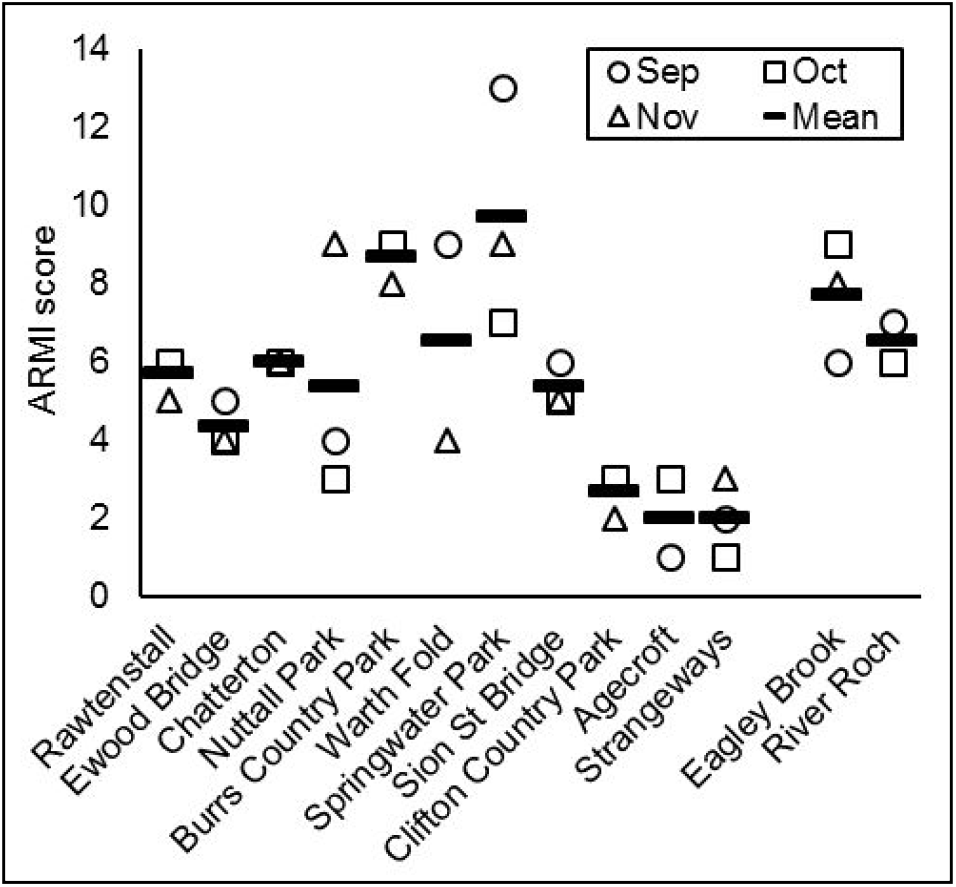
Baseline ARMI scores for 13 sites on the River Irwell and tributaries from autumn 2011

Near continuous long-term monthly ARMI records were compiled from September 2011 for 3 sites (Fig. 3): Ewood Bridge on the upper River Irwell (September 2011 to August 2017; mean 4.41, SD 1.43, range 2-8, n=61); Eagley Brook (August 2011 to August 2017; mean 6.98, SD 1.92, range 4-13, n=58); and the River Roch (November 2011 to May 2015; mean 8.03, SD 2.07, range 3-12, n=37). All three sites show no long-term inter-annual trend. Examining combined annual data for individual months January to December (Fig. 4) showed significant seasonal influence on the ARMI score (P<0.01) using one-way ANOVA.

**Fig. 3.**
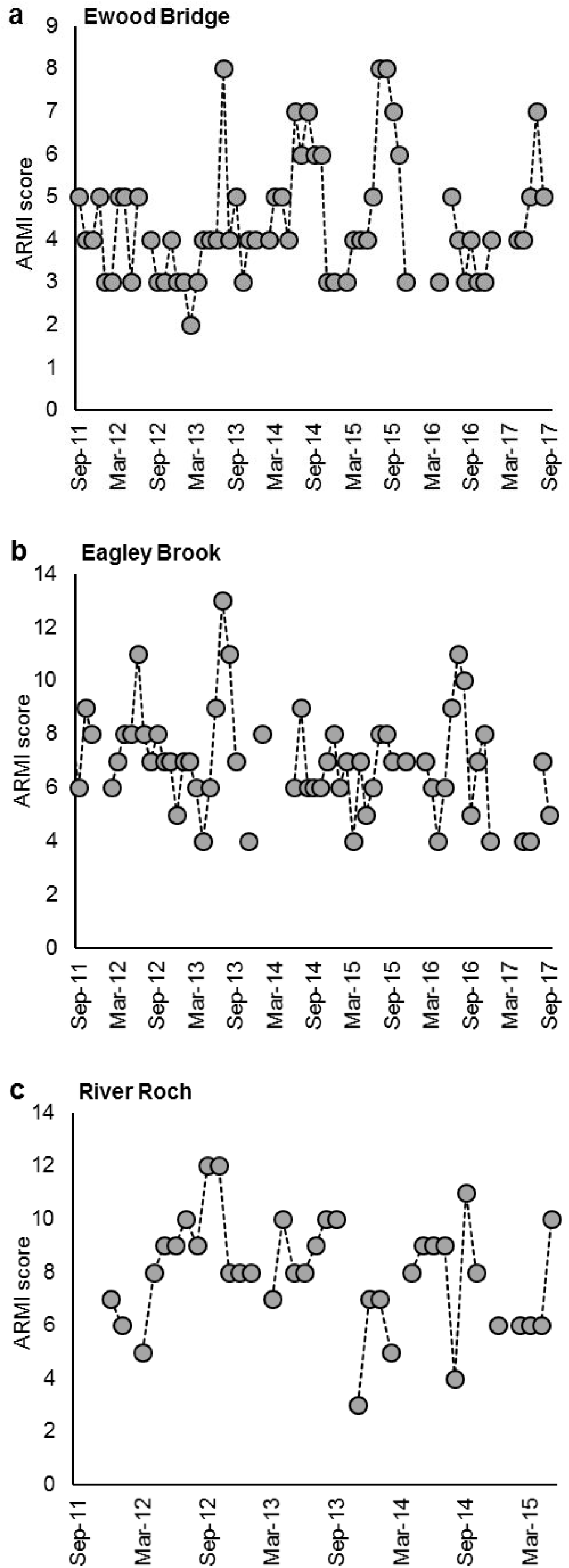
Long-term ARMI scores beginning September 2011 for (a) Ewood Bridge on the River Irwell, (b) Eagley Brook and (c) the River Roch

**Fig. 4.**
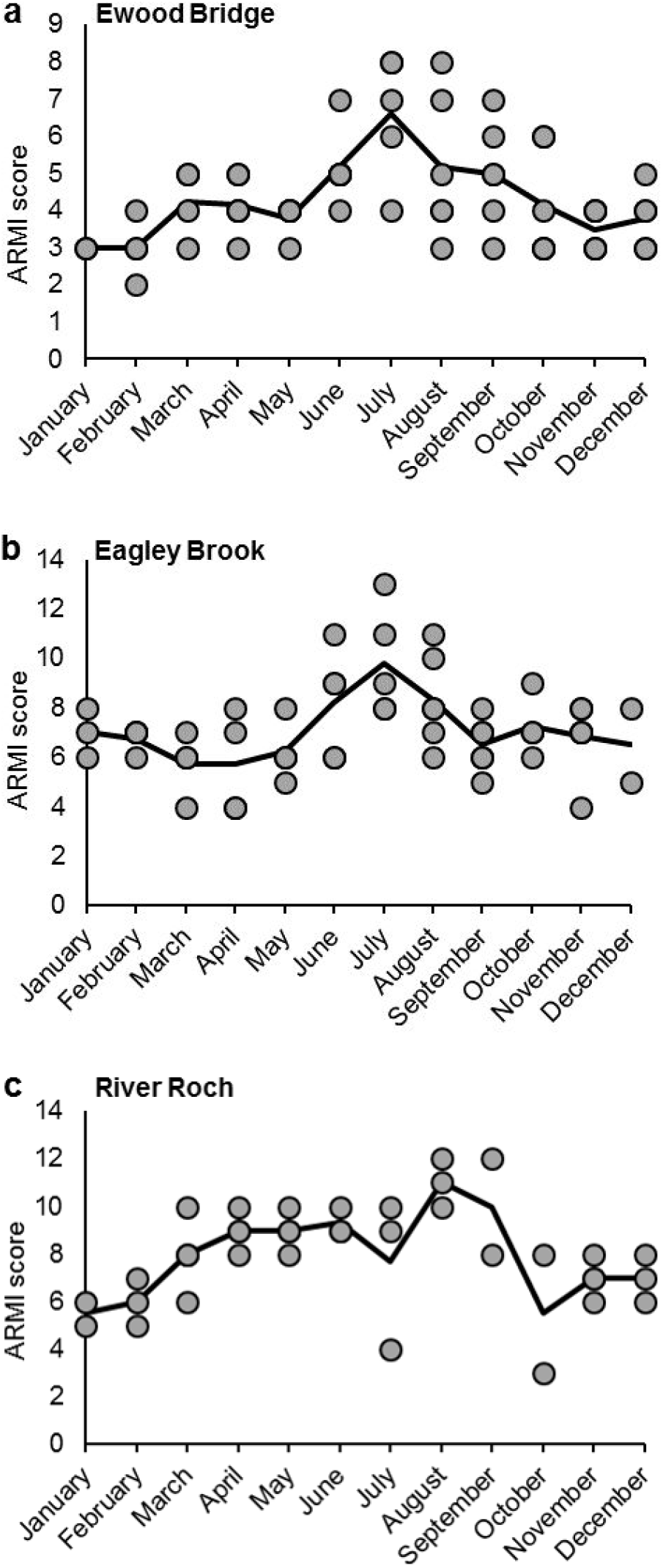
Long-term seasonal deviation of ARMI score by month from the annual mean for (a) Ewood Bridge on the River Irwell, (b) Eagley Brook and (c) the River Roch

The validity of using autumn 2011 ARMI scores as a baseline for the pollution event investigated in April 2017 was therefore tested by comparing multi-year April scores with autumn (September to November scores) for the three long-term monitoring sites. Paired t-tests showed no statistical difference for Ewood Bridge (April mean 4.17, SD 0.75, range 3-5, n=6; autumn mean 4.22, SD 1.31, range 3-7, n=18; n.s.), Eagley Brook (April mean 5.75, SD 2.06, range 4-8, n=4; autumn mean 6.81, SD 1.22, range 4-9, n=16; n.s.) or the River Roch (April mean 9.00, SD 0.82, range 8-10, n=4; autumn mean 7.38, SD 2.50, range 3-12, n=8; n.s.).

### 3.2 Comparison with BMWP and ASPT indices

To compare ARMI, BMWP and ASPT indices, kick samples were taken from 9 sites along the River Irwell between Rawtenstall in the upper catchment and near Manchester city centre at Strangeways (13.5 km to 56.5 km from the source) in January 2012 and then again in March 2012. All macroinvertebrates in each sample were identified and counted and then each individual sample was scored for ARMI, BMWP and ASPT indices. Significant (P<0.01, n=18) moderate positive correlations with ARMI for both BMWP (r^2^=0.653) and ASPT (r^2^=0.653) were assigned using the Spearman correlation coefficient (Fig. 5).

**Fig. 5.**
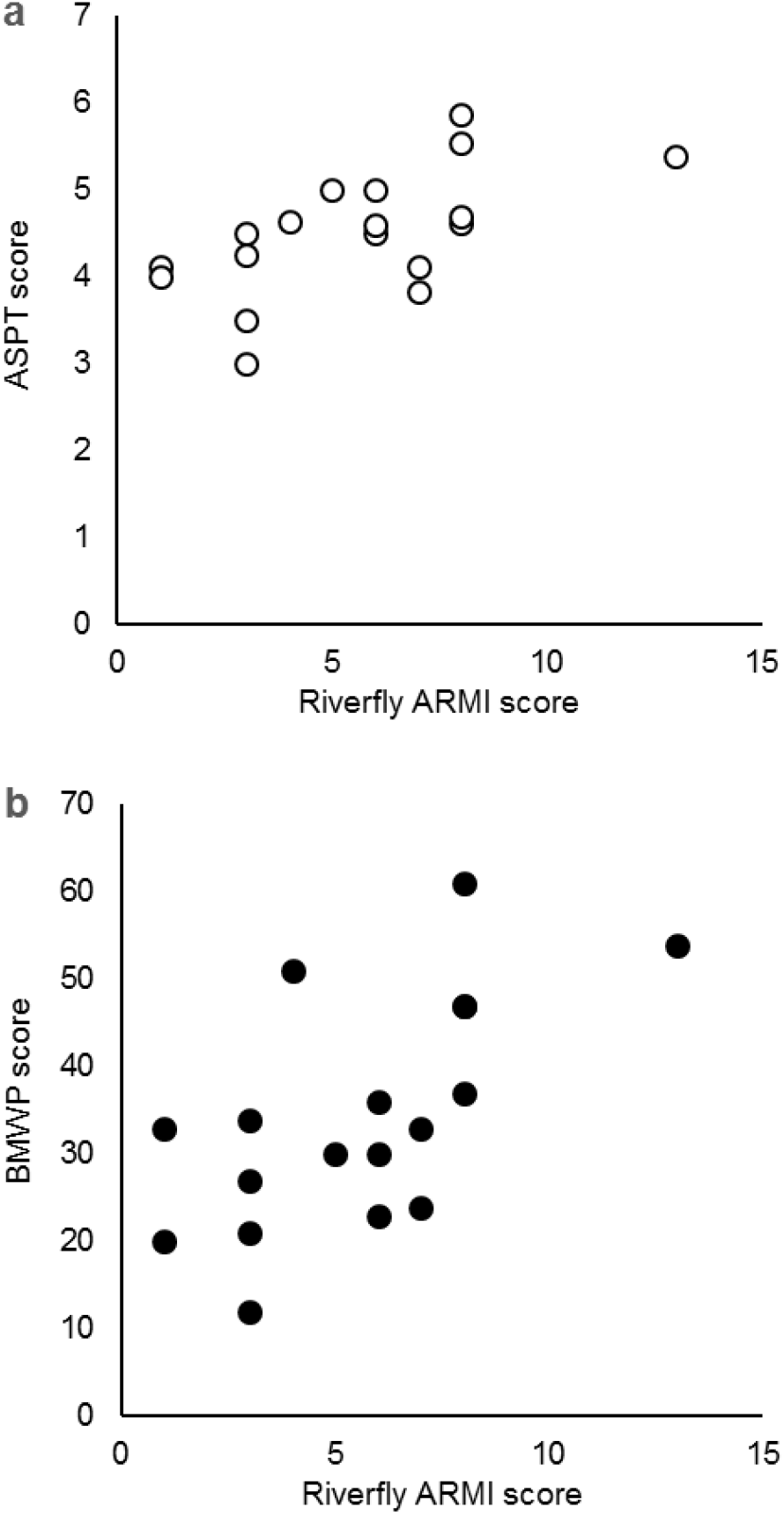
Scatterplot of Riverfly ARMI scores with (a) BMWP and (b) ASPT scores for the same series of samples, taken from 9 sites on the River Irwell in January 2012 and again in March 2012

### 3.3 Macroinvertebrate loss in April 2017

In April 2017 an angler fishing on the River Irwell upriver of Manchester noted large numbers of dead crayfish, suggesting a pollution incident, and contact was made with the SFAS citizen science monitors. A kick sample was carried out to investigate the macroinvertebrate community at the nearest reference site at Burrs Country Park (see Fig. 1) and no macroinvertebrates were found in the sample.

Volunteers carried out further kick samples at existing reference sites upriver to Ewood Bridge, where a healthy macroinvertebrate population was found, and downriver to Agecroft (see Fig. 1) to localize the source of pollution and how far the macroinvertebrate kill extended (see Fig. 6). No macroinvertebrates were found in samples taken at Nuttall Park or Chatterton 6.2 km and 10.0 km upriver of Burrs Country Park respectively but at Ewood Bridge 14.8 km upriver a healthy macroinvertebrate population was found with an ARMI score of 4 typical for the site (long-term mean 4.41, SD 1.43, range 2-8, n=61).

**Fig. 6.**
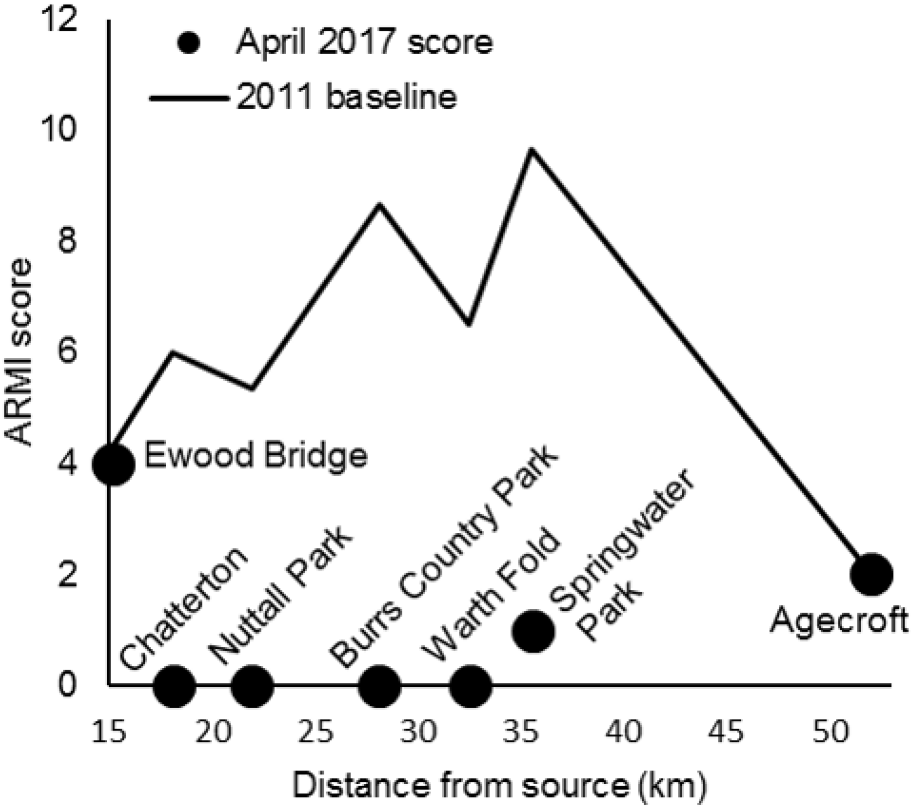
ARMI scores in April 2017 and from the 2011 baseline for 7 reference sites along 37 km of the River Irwell from Ewood Bridge 15 km from the source to Agecroft 52 km from the source

Macroinvertebrates were absent at the first reference site downriver of Burrs Country Park (Warth Fold, 4.4 km along the river). A limited population of macroinvertebrates was found at Springwater Park 7.4 km downriver of Burrs Country Park and 330 m below the confluence of the River Roch; but with an ARMI score of 1 significantly below the baseline of 10 (range 7-13, n=3) recorded from autumn 2011 (P<0.01). Continuing downriver, the third available reference site at Agecroft had an ARMI score of 2, which is comparable to the low baseline score established for this site in autumn 2011 (mean 2, range 1-3, n=3).

### 3.4 Citizen science identification of the putative pollution site with reactive sampling

Volunteers carried out a responsive survey at short distance interval additional sites downriver of the healthy macroinvertebrate population at Ewood Bridge on the River Irwell, a section which includes a discharge point from a wastewater treatment works, and on the River Ogden tributary (see Table 1 and Fig. 7).

**Fig. 7.**
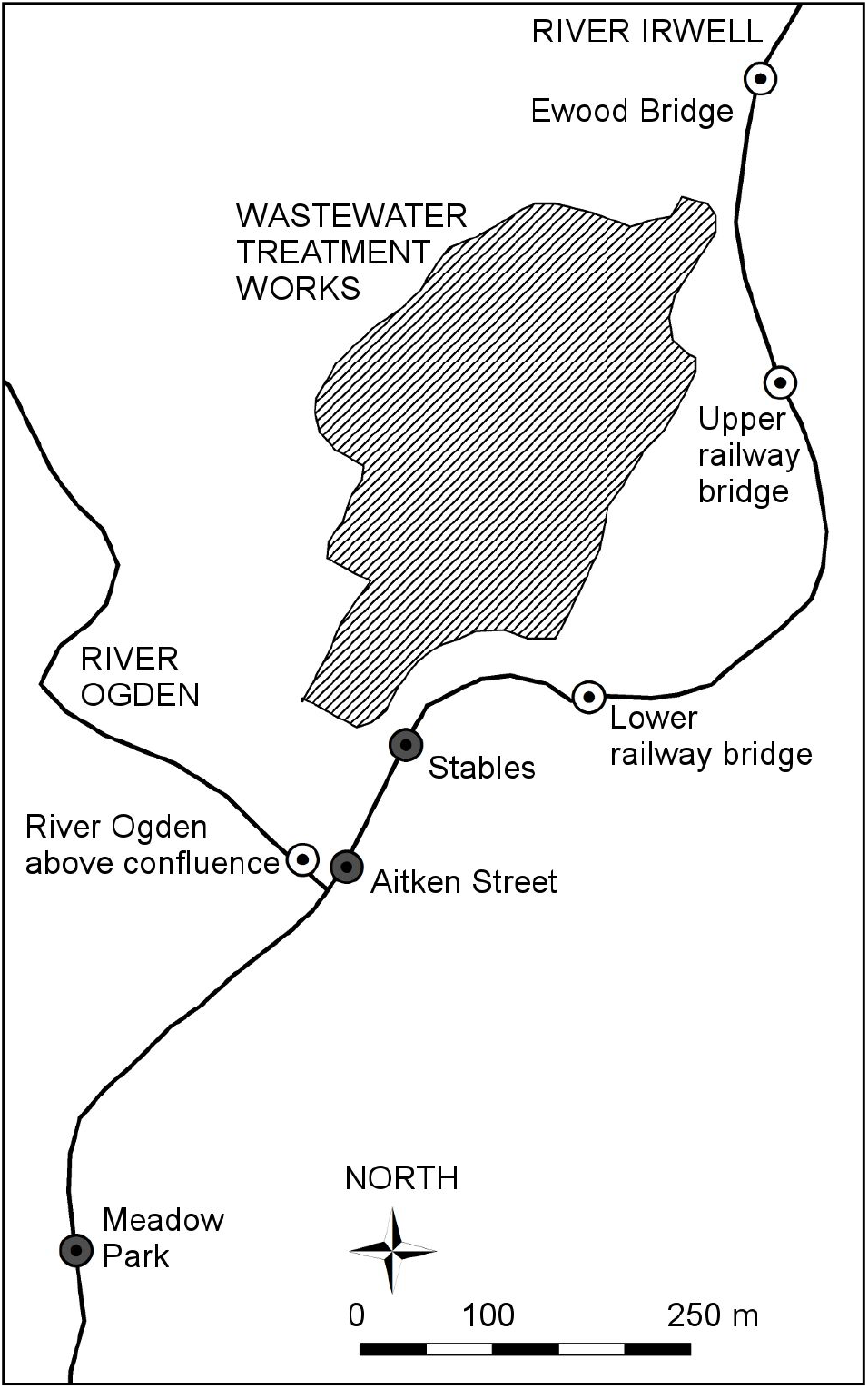
Map of sites sampled downriver of the Ewood Bridge reference site on the River Irwell to identify the stretch within which the April 2017 macroinvertebrate kill began. Grey-filled circles = sites with no macroinvertebrates and white-filled circles = sites with healthy macroinvertebrate samples (ARMI scores of 4 or more)

**Table 1.** ARMI scores for Ewood Bridge and additional sites sampled to identify the stretch of river within which the April 2017 macroinvertebrate kill began

| River and context | Site | ARMI score |
| --- | --- | --- |
| Irwell above wastewater treatment works | Ewood Bridge | 4 |
| Irwell above wastewater treatment works | Upper railway bridge | 4 |
| Irwell above wastewater treatment works | Lower railway bridge | 4 |
| River Ogden | Above confluence with Irwell | 9 |
| Irwell below wastewater treatment works | Stables | 0 |
| Irwell below wastewater treatment works | Aitken Street | 0 |
| Irwell below wastewater treatment works | Meadow Park | 0 |

Healthy macroinvertebrate samples (ARMI score 4) were found upriver of the wastewater treatment works at the two railway bridge crossing sites, with the lower railway bridge approximately 25 m upriver of the discharge point. Macroinvertebrates were, however, absent at the stables site immediately below the wastewater treatment works discharge and at Aitken Street 110 m downriver of the wastewater treatment works discharge. The sample from Meadow Park 490 m downriver of the wastewater treatment works discharge and 380 m downriver of the confluence of the River Ogden also contained no macroinvertebrates. The River Ogden sampled immediately above its confluence with the River Irwell had healthy abundances of various macroinvertebrate groups (ARMI score 9). The location of the start of the macroinvertebrate kill was therefore constrained to an approximately 30 m stretch of the River Irwell alongside the discharge point of the wastewater treatment. The Environment Agency was immediately contacted with the information to request an investigation.

## 4. DISCUSSION

### 4.1 Pollution vigilance: responsiveness of the citizen science network

There will usually be at least one sample taken on the Irwell catchment in any one week as part of the Riverfly Partnership citizen science Anglers’ Riverfly Monitoring Initiative. Two long-term reference sites (Ewood Bridge on the River Irwell and the Eagley Brook tributary) continue to be monitored on a monthly basis, whilst additional volunteers carry out occasional sampling at different ad hoc locations to assess macroinvertebrate populations alongside recreational fishing. Regular monitoring and the presence of anglers ready to take reactive samples if pollution is suspected means large-scale macroinvertebrate kills have a high likelihood of detection. Further investigations by the citizen science group can then be instigated and, if appropriate, be reported to the Environment Agency along with clear data on impacts, as happened in April 2017. Existing reference sites with known baselines allowed volunteers to quickly identify that the April 2017 pollution event had occurred between the Chatterton and Ewood Bridge reference sites. Reactive identification of additional sampling sites was able to constrain a short stretch of river along which the pollution impact began. Reference sites downriver indicated that macroinvertebrate life was absent until the confluence of the River Roch (macroinvertebrates present at Springwater Park 330 m downriver of the confluence, albeit with an ARMI score of 2 significantly below the baseline of 10; P<0.01). This evidence of a substantial pollution event destroying the macroinvertebrate community along an estimated 19 km stretch of the River Irwell was then passed to the Environment Agency requesting an investigatory response. Without the citizen science network provided by the Riverfly Partnership this large-scale macroinvertebrate kill would almost certainly have gone undetected by the Environment Agency because it did not result in evident fish death, which is the usual indicator of a pollution event noticed by the general public. No formal outcome of investigation by the Environment Agency is currently available but the most likely explanation is that a large quantity of pesticide made its way into the River Irwell alongside the wastewater treatment works discharge. This incident has similarities to the chlorpyrifos pesticide pollution events examined by Raven & George (1989) for the River Roding and by Thompson et al. (2016) for the River Kennet, both tributaries of the River Thames in southern England and the latter case similarly identified by citizen scientists following a large-scale macroinvertebrate kill. The Chatterton, Nuttall Park and Burrs Country Park sites with zero macroinvertebrates in April 2017 were monitored for recovery on a monthly basis after the pollution event on the River Irwell. No macroinvertebrates were found in May but olives (Baetidae) were found in small numbers in June and July (10 or less, ARMI score 1-2), increasing to 100 or more olives in September samples (ARMI score 3). No other ARMI macroinvertebrate groups were present as of the latest samples in September 2017.

### 4.2 Broad awareness but limited records

Whilst the ad hoc basis of much of the sampling does suffice to provide a broad awareness of pollution of the rivers of the Irwell catchment and to raise alerts to specific pollution events, it has proven difficult to translate into a resilient long-term monitoring program. The SFAS was only able to engage citizen science volunteers to continue monitoring for 5 of the initial 13 sites used in the autumn 2011 baseline survey. Of these 5, monitoring ended at Rawtenstall after 20 months (data last collected April 2013) and at Clifton Country Park after 23 months (data last collected June 2013). Monitoring continued on the 3 remaining sites, giving long-term records, and continues as of September 2017 for both Ewood Bridge on the upper River Irwell and on the Eagley Brook tributary (monitoring ended on the River Roch in May 2015). Despite additional training events in 2015, 2016 and 2017 certifying further volunteers as monitors, the informal and devolved organization of SFAS citizen science activity has hampered efforts to restart long-term monitoring at other reference sites. Two of the 2016 trainees do regularly conduct Riverfly Partnership assessments of ad hoc sites alongside recreational fishing, as do other interested anglers, but it has proven difficult to capture this data and the volunteer group’s central database is far from comprehensive. The Riverfly Partnership does have an online portal for record submission but this only allows data entry for monitoring sites formally agreed with the Environment Agency, meaning there is no flexibility for additional data capture. Approaches to sustaining volunteer engagement should be reappraised considering the importance of maintaining links between the data collectors, the accumulating data, and the monitoring outcomes as stressed by Roy et al. (2012) and Pocock et al. (2014). One possibility to establish durable monitoring sites might be to engage a coordinated network of existing local environmental volunteer groups active along rivers, for example associated with parks, woodlands and other green spaces.

### 4.3 Approaches for an effective citizen science monitoring network across Greater Manchester

A major pollution event such as occurred in April 2017 needs to be detected with minimal delay so that action can be taken to remedy the source and for the statutory authorities to obtain further investigatory evidence. Realistically, monitoring activity once a week is the maximum temporal resolution for even the most vigorous volunteer group. Yet any one specific site cannot be sampled at greater than monthly frequency because of the need to allow recolonization and normalization of the habitats in between sampling events. In addition, a point source pollution event will only be detected if a sample is taken from a site downriver. An effective monitoring network therefore depends on appropriate coverage across the Irwell catchment and a coordinated roster of sites for weekly samples. This would offer the opportunity to coordinate the three goals competing for the finite time of citizen science monitors: frequent monitoring able to detect major pollution events with minimum delay; breadth of coverage; and specific sites with long-term records. This coverage and frequency is only possible with volunteers (Latimore & Steen 2014; Kobori et al. 2016) but also depends on individual monitors being effectively supported, which has proven a time and resource challenge within the volunteer-managed SFAS citizen science program.

### 4.4 Deciding pollution alert trigger levels in urban river catchments with low and variable macroinvertebrate diversity

Unless a pollution event causing large-scale macroinvertebrate death also results in a major fish kill it is likely to be unseen, unreported and hence not investigated without ARMI activity. The ARMI methodology was designed specifically to enable trained citizen scientists to detect severe changes in water quality such as this. However, a challenge has been in justifying an investigatory response by the Environment Agency to low ARMI scores, particularly when these are only localized drops in macroinvertebrate abundance. Based on 6 years data, for example, Ewood Bridge has a mean ARMI score of 4.41 (SD 1.43, range 2-8) so an ARMI score of 1 is the realistic trigger level and an investigation by the Environment Agency would only be triggered in response to a major macroinvertebrate kill. The importance of a baseline for specific sites is highlighted by the baseline drop, for example, between Springwater Park (ARMI score 10) and Agecroft (ARMI score 2) 16.5 km downriver. The April 2017 ARMI score of 2 for Springwater Park indicates an impacted macroinvertebrate community, whereas the ARMI score of 2 for Agecroft is consistent with the baseline for this site closer to Manchester city centre. Whether that means the pollution impact of the April 2017 event was negligible at Agecroft we are unable to judge, given the lack of sensitivity of such a low baseline and already markedly degraded community.

### 4.5 The Anglers’ Riverfly Monitoring Initiative as a practical and effective citizen science method

For use in citizen science, a sampling method must be easy to use and require minimal equipment and training. Kick sampling, the semi-quantitative approach of both Riverfly Partnership citizen science and UK statutory monitoring, is favored for statutory biomonitoring globally over complete and quantitative sampling methods such as Surber sampling because it is quick and easy to carry out (Everall et al. 2017). Field sorting becomes practical when using live samples and identification to broad taxonomic groups without optical aids (Haase et al. 2004; Growns et al. 2016). The Riverfly Partnership ARMI satisfies these requirements through simple bankside identification of live samples with broad groups of readily identifiable taxa and minimal equipment requirements. Groups are identified according to presence and number of tails and pairs of legs, appearance of the gills, and (for cased caddisfly larvae) presence of a case. Live samples make identification easier through behavioral characteristics – the only two groups that look superficially similar as dead specimens are readily distinguished in a live sample by their very different swimming motions (i.e. the short forward dashes of olives Baetidae versus the convulsive thrashing of blue-winged olive Ephemerellidae). Experienced monitors can score a kick sample spread out in a small tray by eye for the ARMI index without any sorting needed. Even for beginners, scoring for abundance is straightforward after separating the 8 target groups by live sorting into a compartmentalized tray (although this is relatively time consuming). Exact counts are difficult with moving live individuals but estimates of numbers suffice for the Riverfly Partnership ARMI abundance categories. Broad taxonomic groups are recognized as making little difference to biotic indices scoring compared to species-level identification and accepted as the only option for citizen science monitoring where taxonomic expertise, time, and equipment are limited (Hilsenhoff 1988; Lenat & Resh 2001; Jones 2008). The rapid response and immediate results obtained at the time of sampling is a key advantage of the Riverfly Partnership ARMI index over the BMWP and ASPT indices. The potential for geographic coverage and temporal resolution that the ARMI index gives would be unattainable using more discriminatory scoring methods such as BMWP or ASPT, a major advantage of citizen science approaches (Kobori et al. 2016; Shupe 2016). Comparison here of ARMI scores with BMWP and ASPT scores for the same samples found statistically highly significant (P<0.01) correlations, validating the ARMI as an equivalent indicator of water quality status for the rivers of the Irwell catchment. A larger data set, however, would be needed to derive quantitative relationships that could be applied more widely – this should cover the full range of ARMI scores, seasons and site types including substrate and flow regime (the latter influenced by canalization in urbanized rivers such as the Irwell). One important aspect of the ARMI is that it assesses a selection of taxa found in clean to moderately polluted water only, rather than all macroinvertebrates including those that can live in highly polluted water as with the BMWP and ASPT indices. The ARMI therefore would not be sensitive to differences between highly and severely polluted water as scores would simply be zero – whilst BMWP and ASPT indices retain sensitivity within more polluted water. It is also interesting to note that, despite anecdotal reports from anglers of less macroinvertebrate life seen when fishing, the extensive winter floods of 2015-16 across northern Britain (Marsh et al. 2016) had no discernible impact on ARMI scores in the monitoring records for the two Irwell catchment sites which cover this period (Ewood Bridge and Eagley Brook). This could be because broad identification groups miss subtler changes (Lenat & Resh 2001). It certainly highlights that the variability of ARMI scores means only severe pollution impacts on macroinvertebrate life are identifiable in the Irwell catchment. The ARMI index was of course explicitly designed to detect acute pollution incidents but this highlights that more detailed monitoring and investigation work would be required to identify subtler chronic pollution and to assess its impacts. In relation to these observations, pilot studies are ongoing within the Riverfly Partnership for “Riverfly Plus” indices of greater complexity that would provide citizen scientists with a monitoring platform sensitive to specific water quality stressors beyond the scope of the existing ARMI index.

### 4.6 Citizen science as a catalyst for public engagement and change

The involvement of volunteers and local communities allows monitoring to transcend the limitations of research and monitoring by academics or regulatory bodies and provides scope for implementation of community-led environmental improvement projects (Kobori et al. 2016). This is very much the case with the Riverfly Partnership citizen science program in Greater Manchester. The Riverfly Partnership group at the SFAS galvanized interest from anglers in wider environmental action. This led to further projects and funding from various sources for tackling invasive non-native species such as Giant Hogweed, Himalayan Balsam and Japanese Knotweed along the Irwell and tributaries. The SFAS also became involved in community clean up events along the riverside in public green spaces and in schools outreach engaging children with their local rivers. This community engagement led to media interest, which raised the profile of events and encouraged more people to get involved. As was typical for friendly societies (Cordery 1995), the SFAS has held its regular meetings in public houses for 200 years and this history led to the group and the citizen science work being featured on the British Broadcasting Corporation (BBC) television series “Pubs That Built Britain” in April 2015. The BBC’s Countryfile magazine has also featured the SFAS’ citizen science, wider environmental work and schools outreach (Griffiths 2016). As has been observed elsewhere (Nerbonne & Nelson 2008), conflicts of volunteer goals, however, were quickly apparent with some anglers not wanting the group to move in an environmental direction and non-anglers wanting to get involved but being discouraged by the SFAS being an “anglers’ society”. The divergence of goals and interest outside the SFAS led to the establishment in 2015 of the Mersey Basin Rivers Trust as a free membership umbrella organization that would be the face of the wider environmental work.

### 4.7 Bridging the gap between top-down initiatives and communities

Being part of the wider River Mersey catchment, the Irwell catchment benefited substantially from the Mersey Basin Campaign between 1985 and its termination in 2010. This was a ground-breaking model of partnership approach linking water quality improvements with wider river corridor sustainability and urban regeneration (Struthers 1997; Wood et al. 1999). Government-led but with substantial private investment, such partnerships included the regeneration of Manchester Docks on the River Irwell (where it becomes canalized as the Manchester Ship Canal) and its transformation into the Salford Quays development (Williams et al. 2010). The Healthy Waterways Trust was the Mersey Basin Campaign vehicle for administering charitable funds and continued post-2010 as the focus for continuing government-led input and resources for engaging people with the River Irwell and the wider Mersey Basin, later rebranding as the Healthy Rivers Trust. Yet there remained a need for more effective community engagement, with criticism of the top-down approach for genuinely engaging communities (Eden & Tunstall 2006). The expansion of SFAS and Riverfly Partnership citizen science into wider environmental work and the foundation in 2015 of the Mersey Basin Rivers Trust went someway to filling this gap. The Mersey Basin Rivers Trust has now merged with the Healthy Rivers Trust, bridging the gap between top-down and bottom-up approaches. The new entity, the Mersey Rivers Trust, has taken the lead role in both the Upper Mersey (covering the Irwell and tributaries) and Lower Mersey catchment partnerships under the UK government’s catchment-based approach (CaBA) framework for EU Water Framework Directive compliant river basin management plans (Cascade Consulting, 2013; Robins et al. 2017).

## 5. Conclusion

The Riverfly Partnership citizen science network provides an important regular presence on riverbanks across the United Kingdom identifying pollution incidents that would otherwise likely go unseen and unreported. Anglers trained as citizen scientists provide day-to-day coverage from the riverbanks whilst fishing that cannot be matched for awareness by cost- and labor-intensive professional assessments of water quality. Riverfly Partnership citizen science in Manchester has catalyzed broader environmental engagement, further funding, and led to the bridging of a divide between top-down government-led approaches and genuine grassroots environmental action. The ability of citizen science networks such as this to provide pollution vigilance across a river catchment was demonstrated in April 2017 by detecting a major pollution event and rapidly constraining the location of the pollution source. Challenges remain in terms of coordination, centralized record-keeping and reconciling different goals with limited resources but this case study shows the potential of citizen science to play an important role in the protection and improvement of river catchments.

## Acknowledgements

With thanks to all the people who have contributed their time, skills and resources to macroinvertebrate monitoring, community events and wider environmental work at the Salford Friendly Anglers’ Society. Particular thanks to Mike France, Tony Quinn, Arthur Hamer, Dave Vickery and Ian Hedley for their collection of monthly monitoring data and for insights and to Tara Barbour who collected the BMWP and ASPT data.

